# Vesicular GABA-transporter neurons in the zona incerta are maximally active during non rapid-eye movement (NREM) and rapid-eye movement (REM) sleep

**DOI:** 10.1101/2020.04.15.043372

**Authors:** Carlos Blanco-Centurion, SiWei Luo, Aurelio Vidal-Ortiz, Priyattam J. Shiromani

## Abstract

Sleep and wake are opposing behavioral states controlled by the activity of specific neurons. The neurons responsible for sleep/wake control have not been fully identifed due to the lack of *in-vivo* high throughput technology. We use the deep-brain calcium (Ca^2+^) imaging method to identify activity of hypothalamic neurons expressing the vesicular GABA transporter (vGAT), a marker of GABAergic neurons. vGAT-cre mice (n=5) were microinjected with rAAV-FLEX-GCaMP6_M_ into the lateral hypothalamus and 21d later the Ca^2+^ influx in vGAT neurons (n=372) was recorded in freely-behaving mice during waking (W), NREM and REM sleep. Post-mortem analysis revealed the lens tip located in the zona incerta/lateral hypothalamus (ZI-LH) and the change in fluorescence of neurons in the field of view was as follows: 54.9% of the vGAT neurons had peak fluorescence during REM sleep (REM-max), 17.2% were NREM-max, 22.8% were wake-max while 5.1% were both wake+REM max. Thus, three quarters of the recorded vGAT neurons in the ZI-LH were most active during sleep. In the NREM-max group Ca^2+^ fluorescence anticipated the initiation of NREM sleep onset and remained high throughout sleep (NREM and REM sleep). In the REM-max neurons Ca2+fluorescence increased before the onset of REM sleep and stayed elevated during the episode. Activation of the vGAT NREM-max neurons in the zona incerta and dorsal lateral hypothalamus would inhibit the arousal neurons to initiate and maintain sleep.

## Introduction

A distributed network of neurons is hypothesized to regulate the alternation between waking, non-REM sleep and REM sleep. Some neurons, such as those containing norepinephrine (NE), serotonin (5-HT), histamine (HA) or orexin, are most active during waking, decrease their activity in NREM sleep, and are silent in REM sleep [reviewed by (Jones, 2020). Specific manipulations that increase or decrease activity of these neurons have a predictable effect on waking levels indicating that these neurons induce and maintain the waking state. However, what phenotype of neurons initiates and maintains sleep? Such a neuronal phenotype is likely to be inhibitory and clusters of GABAergic neurons in the preoptic (Chung et al., 2017) and lateral hypothalamic (Hassani et al., 2010) areas that discharge maximally in NREM sleep and REM sleep have been found. The discovery of sleep active GABAergic neurons is not surprising because GABA is the primary inhibitory neurotransmitter in the CNS. Norepinephrine, serotonin, histamine and orexin neurons are all wake-active (Jones, 2020). However, are all GABAergic neurons sleep-active or do they have a heterogenous pattern of activity associated with waking, NREM and REM sleep? It is crucial to answer this question because then specific neurons can be manipulated to predictably regulate sleep and wake states, especially disorders of sleep such as insomnia (too little sleep) or narcolepsy (excessive sleep). The *in-vivo* electrophysiology method is the primary method that has been used to measure activity of neurons during waking, NREM and REM sleep (Aston-Jones and Bloom, 1981; John et al., 2004; Wu et al., 2004; Lee et al., 2005; Mileykovskiy et al., 2005). However, there are limitations to this method and new recording tools are required to rapidly disentangle the phenotype of neurons underlying behaviors such as sleep.

A new method that can image neuronal activity was recently introduced (Ghosh et al., 2011). This method measures calcium (Ca^2+^) intracellular gradients associated with excitatory signaling in neurons (Tian et al., 2012). A genetically encoded Ca^2+^ fluorescent indicator (GCaMP6) expressed in specific neurons provides a reliable readout of the neuron’s activity level (Chen et al., 2013). The change in Ca^2+^ induced fluorescence in GCaMP6 expressing neurons is recorded via a miniature single-photon epifluorescence microscope attached to a miniature gradient index (GRIN) lens (Ghosh et al., 2011). Deep-brain Ca^2+^ imaging has proven to be very useful in confirming existing *in-vivo* electrophysiology data by imaging neurons in quiet waking, NREM and REM sleep (Weber et al., 2015; Cox et al., 2016; Weber and Dan, 2016; Chung et al., 2017; Weber et al., 2018). We recently used the deep-brain Ca^2+^ imaging method to monitor activity in neurons expressing melanin concentrating hormone (Blanco-Centurion et al., 2019). Here we use the deep-brain Ca^2+^ imaging method to monitor activity of neurons that constitutively express the gene for vesicular GABA transporter (vGAT) gene, and thus, synthesize and release GABA.

In the posterior hypothalamus, GABAergic neurons are intermingled with the orexin and histamine neurons that promote arousal and are closely mingled with MCH neurons that, in contrast, promote sleep. However, the vGAT neurons are separate and distinct from the orexin, histamine or MCH neurons, and a subset of vGAT neurons show increased Ca^2+^ fluorescence during appetitive or consummatory behavior (Jennings et al., 2015). In the current study we find that approximately three-quarters of the vGAT neurons located in the zona incerta-dorsal lateral hypothalamus were maximally active in both NREM and REM sleep. Thus, unlike the arousal neurons which are all active in waking but silent in sleep, we find that the activity of GABA neurons is heavily biased towards sleep. Activation of the vGAT neurons in the ZI-dLH would inhibit the arousal neurons enabling the brain to fall asleep.

## Methods

### Animals and surgery

The VA Institutional Animal Care and Use Committee approved all animal use procedures (protocols # 638,639). The vGAT-IRES-Cre (Jackson Laboratories, JAX stock #016962) mice were bred in our animal facility and those found to be Cre+ (Transnetyx, Inc.) were later chosen for the imaging study. After surgery, the animals were singly housed in polycarbonate cages with α-cellulose bedding. Standard rodent laboratory food pellets and water were available *ad-libitum*. The temperature was controlled (25°C± 2) while a 12:12h light-dark cycle (6AM-6PM lights on; 300 lux) was always maintained. The mice used for this study were four months old and had a mean body weight of 34.7 g [±1.71g] at the time of surgery (Table 1).

**Table 1.** Sex, age, body mass and numbers of neurons recorded in the study.

| Mouse ID | Sex | Age (month) | Body weight (g) | Numbers of vGAT neurons that were imaged |
| --- | --- | --- | --- | --- |
| vG176058 | M | 4 | 36.4 | 22 |
| vG194239 | M | 4 | 39.4 | 93 |
| vG194243 | M | 4 | 28.9 | 77 |
| vG194247 | F | 4 | 34.5 | 108 |
| vG194251 | F | 4 | 34.3 | 72 |
| Average (+/- SEM) |  | 4 | 34.7 (1.71) | 78.4 (11.1) |
| **Total** | **60%M** |  |  | 372 |

The mice were deeply anesthetized (2% isoflurane) and, using a stereotaxic apparatus, the genetically encoded Ca^2+^ indicator, GCaMP6m (rAAV-DJ-EF1a-DIO-GCaMP6m, titer: 0.4 ×10^9^/ml GC, vol:2-4 µl: Stanford University Vector Core), was slowly injected over a 10 min period into the lateral hypothalamus (relative to Bregma A=-1.6; lateral=0.7; ventral from brain surface=4.8mm). The viral injection cannula was held in place for 45 minutes and then slowly retracted. After delivery of the virus, a GRIN lens (0.6mm diameter; 7.3mm length; Inscopix stock number 1050-002208) was inserted into the same cannula track 0.1 mm above the viral transfection target. At this time four miniature screw-type electrodes (Plastic One Inc) were secured to the skull to record the electroencephalogram (EEG) activity. To measure the electromyogram (EMG) activity, two multi-stranded wire electrodes were inserted into the nuchal muscles. The EEG, EMG electrodes and the GRIN lens were permanently attached to the skull with dental cement. For pain management, mice were administered carprofen (5 mg/kg sc) for two days. Ten days later, mice were anesthetized (2% isofluorane), placed in the stereotaxic instrument, and using a micromanipulator (Sutter Instruments) a single photon miniature fluorescence microscope (Inscopix nVista) attached to a baseplate was carefully positioned atop the GRIN lens. Once the best focal plane was achieved, the baseplate was carefully cemented to the headset. The miniscope was detached and the mice returned to the vivarium.

Twenty-one days after the injection of the AAV the miniscope and the sleep recording cable were attached and the mice were adapted to the recording environment for 6-8 h during the day cycle. The adaptation took place for 3 successive days. At the end of each adaptation period the miniscope was detached, a protective cover plate was attached to the baseplate and mice were returned to the vivarium.

### Acquisition of the Ca^2+^ imaging and sleep data

Sleep and Ca2+imaging was done during the second half of the lights-on period. The first half was reserved for adapting the mice to the recording environment. Ca^2+^ imaging data were recorded with the nVista^®^ software (Inscopix, Inc) at a sample rate of 10 frames per second, camera gain of 3.0X and under 10% LED blue radiance power (0.1mW at the GRIN lens bottom). Each mouse we always imaged at a single focal plane that gave us the sharpest images of the fluorescent neurons. EEG/EMG raw analogue signals were bandpass filtered (EEG:0.1-60Hz; EMG:100-1KHz) and amplified by Grass 7P51 amplifiers, and then digitized with the OmniPlex^®^ acquisition system (Plexon, Inc). OmniPlex also recorded video of the animal’s behavior (CinePlex Studio^®^). The start signal from the miniscope was input to OmniPlex. Thus, OmniPlex displayed the acquired signals from five distinct channels: two EEG, one EMG, one nVista recording signal and one from the video camera. For each mouse the data were acquired for a mean of 50.2min [±2.4min] which allowed sampling of several cycles of wake, NREM and REM sleep.

### Ca^2+^ imaging data offline processing

Imaging data were processed and analyzed in an independent and blind manner. Raw imaging data were processed using the Inscopix Data Processing software (Inscopix DPS^®^, Inc.). For each mouse raw nVista HDF5 imaging files were temporally reduced by a factor of 10 (i.e.10Hz down 1 Hz) and corrected for defective pixilation, row noise and dropped frames. The downsampled movie was then spatially bandpass filtered (0.005-0.5 pixels)and then repeatedly corrected for motion artifacts until it rendered the steadiest calcium signal achievable (<3 µm of either X, Y axis translation). Subsequently, the movie was processed with the *mean projection image* tool that yielded a single reference frame (F_0_). The reference frame (F_0_) represents the mean Ca^2+^ fluorescence intensity across the recording period and independently of the sleep states. The change in fluorescence (ΔF) is (F*i*-F_0_)/F_0_) where *i* represents each movie frame, and were automatically extracted by a tandem run of Principal and Independent Component Analysis (PCA-ICA) algorithm. After the PCA-ICA extracted the *potential* neurons (regions of interest), each probable neuron was manually vetted (accepted or rejected) to determine that it showed a peak Ca^2+^ trace featuring a rapid rise followed by a slow decay and the *maximum projection image* showed neuronal soma-like morphology with an appropriate size (10-20 µm).

### Sleep and behavioral data offline processing

The sleep-wake states were identified based on standardized criteria for EEG/EMG activity patterns and video recordings of the behavior of the mouse (CinePlex Editor^®^; Plexon.com). Sleep scoring was done manually using specialized software (Neuroexplorer.com). Active waking (AW) was identified when mice were behaviorally active (grooming, walking, rearing) with desynchronized EEG and high EMG activity. Quiet waking (QW) was identified by the occurrence of desynchronized EEG, lower EMG tone relative to active waking, and by lack of behavioral activity. Non-REM (NREMS) sleep consisted of high amplitude, slow frequency waves (0.5-4 Hz) in the EEG together with a low EMG tone relative to waking and a sleeping posture displayed on the video. REM sleep was identified by the presence of regular EEG theta activity (4-8 Hz) coupled with low EMG relative to NREMS or waking. During REM sleep the mouse exhibited a relaxed sleeping posture. The video camera’s resolution and field of view made it difficult to detect the phasic aspects of REM sleep such as muscle twitches or vibrissae movements. However, during REM sleep mice displayed an irregular and fast breathing pattern that is different from the breathing pattern seen in NREMS. Lastly, we characterized a REM sleep transitional state (Rt) which represented the 15s period before onset of the REM sleep episode. REM sleep transition is characterized by a mix of NREMS and REM sleep EEG and EMG activity. Sleep scoring was always done by a person blind to the Ca^2+^ imaging data.

### Experimental Design and Statistical analysis

A generalized linear mixed model (SPSS-25) with post-hoc Bonferroni paired comparisons (sequential adjusted) was chosen to determine statistical significance. An unstructured covariance matrix yielded a low Akaike Information Criteria and the best model fit. The GLMM model was appropriate based on the correlation of the fluorescence data between the neurons, the skewed distribution of the data (Kolmogorov-Smirnov test for normality), and unbalanced design. Statistical significance was evaluated at the P=0.05 level (two-tailed). Graphs were plotted using GraphPad Prism.

### Post-mortem analysis of GCaMP6m expression

After Ca^2+^ imaging was completed, mice were euthanized with 5% isofluorane and then perfused transcardially with 0.9% saline (10 ml) followed by 10% buffered formalin in 0.1M PBS (30ml). The brains were collected and fixed in 10% buffered formalin for one week. The brains were equilibrated in 30% sucrose solution in 0.1 M PBS and cut coronally at 40 microns. The sections were washed and mounted onto gelatin coated slides and cover slipped using fluorescence mounting medium. The fluorescence images were obtained at 10x and 20x using Leica TCS-SP8 confocal microscope. GCaMP6m positive neurons were directly identified on the specimens because transfected vGAT-cre neurons specifically expressed the circularly permuted EGFP (Nagai et al., 2001).

### Data and material availability

All raw and extracted data, code and materials used in the study are available by writing to the corresponding authors. We encourage data mining of the imaging data set to extend the analysis.

## Results

### Location of the GRIN lenses and the vGAT transfected neurons

Figure 1 depicts a representative image of the GRIN lens atop a cluster of transfected vGAT neurons. In all mice the transfected vGAT neurons were located most densely within the ventral zona incerta (vZI) with few neurons below the ZI in the dorsal lateral hypothalamus (dLH). We note that this is the same region containing MCH neurons (Blanco-Centurion et al., 2019). The tip of the GRIN lens ended in the zona incerta as depicted in Figure 1. We noticed that a virus titer of 4×10^8^ GC/ml gave us high numbers of transfected vGAT neurons. The neurons appeared to be very healthy with the GCaMP6m expression inside the somata and absent from the cell’s nucleus. Because of the dense cluster of vGAT neurons in the vZI area, we were able to image multiple neurons within a single field of view (Panel C; fig. 1).

**Figure 1.**
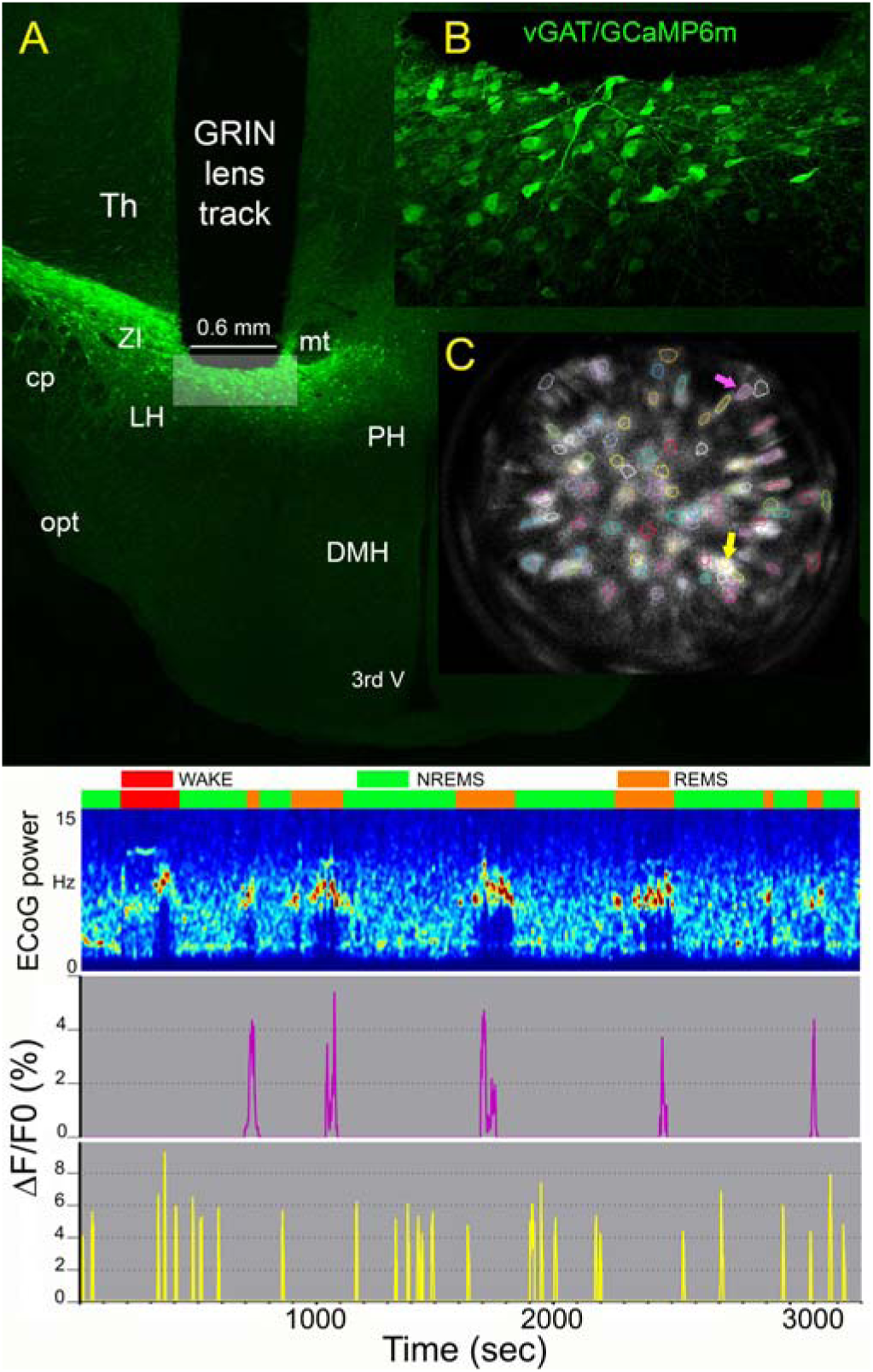
Expression of the Ca^2+^ indicator GCaMP6m in vGAT-cre neurons in the zona incerta and dorsal lateral hypothalamus (ZI-dLH). Panela A depicts the location of the GRIN lens with the tip in a dense cluster of vGAT neurons in the zona incerta. Inset B is a closeup of the embossed rectangle in panel A and shows numerous vGAT-GCaMP6 containing neurons within the focal plane of the GRIN lens. Panel C illustrates the field of view of the GRIN lens with the region of interests extracted by PCA-ICA analysis. PCA-ICA also calculated the neurons’ contours (colored circles). Contours indicate regions of interest where co-variation of fluorescent signal was found to be significantly above the mean projection image. Panel D depicts a one-hour recording of the sleep-wake states (colored boxes) in one mouse. The change in cortical EEG during wake, NREM and REM sleep is depicted as a heat map (0-15Hz). The change in fluorescence in two neurons (represented by two arrows in panel C) is depicted in panel E. Purple trace shows that the change in fluorescence in one neuron is associated with REM sleep while the yellow trace illustrates a different neuron with activity that is linked to NREM sleep. *Abbreviations*; DMH=Dorsomedial hypothalamus, LH=lateral hypothalamus, PH=Posterior hypothalamus, Th=Thalamus, ZI=Zona incerta; cp=cerebellar peduncle, mt=mammillothalamic tract, opt=optical tract, 3rdV=third ventricle.

### Activity of vGAT neurons in waking and sleep

Cytoplasmic Ca^2+^ fluorescence in vGAT neurons was recorded during sleep states and during periods of quiet or active waking. Periods of walking, rearing, or grooming were classified as active waking. A total of 372 individual neurons were imaged in five vGAT-cre mice (Table 1).

The activity of the 372 vGAT neurons was binned into four groups (Figure 2) based on clear and precise criteria. Neurons that were most active during REM sleep (REM-max group) represented most of the imaged neurons or 54.9% of the total (n=204). The general linear mixed model found that the average Ca^2+^ fluorescence during REM sleep was significantly higher compared to waking, NREM or REM-transition states (F[4,965]=30.915; P=0.0001). The fluorescence in REM sleep was roughly eight times as high as it was measured during the other vigilance states. REM sleep-max neurons also showed mild fluorescence during active waking or during the transition into REM sleep.

**Figure 2.**
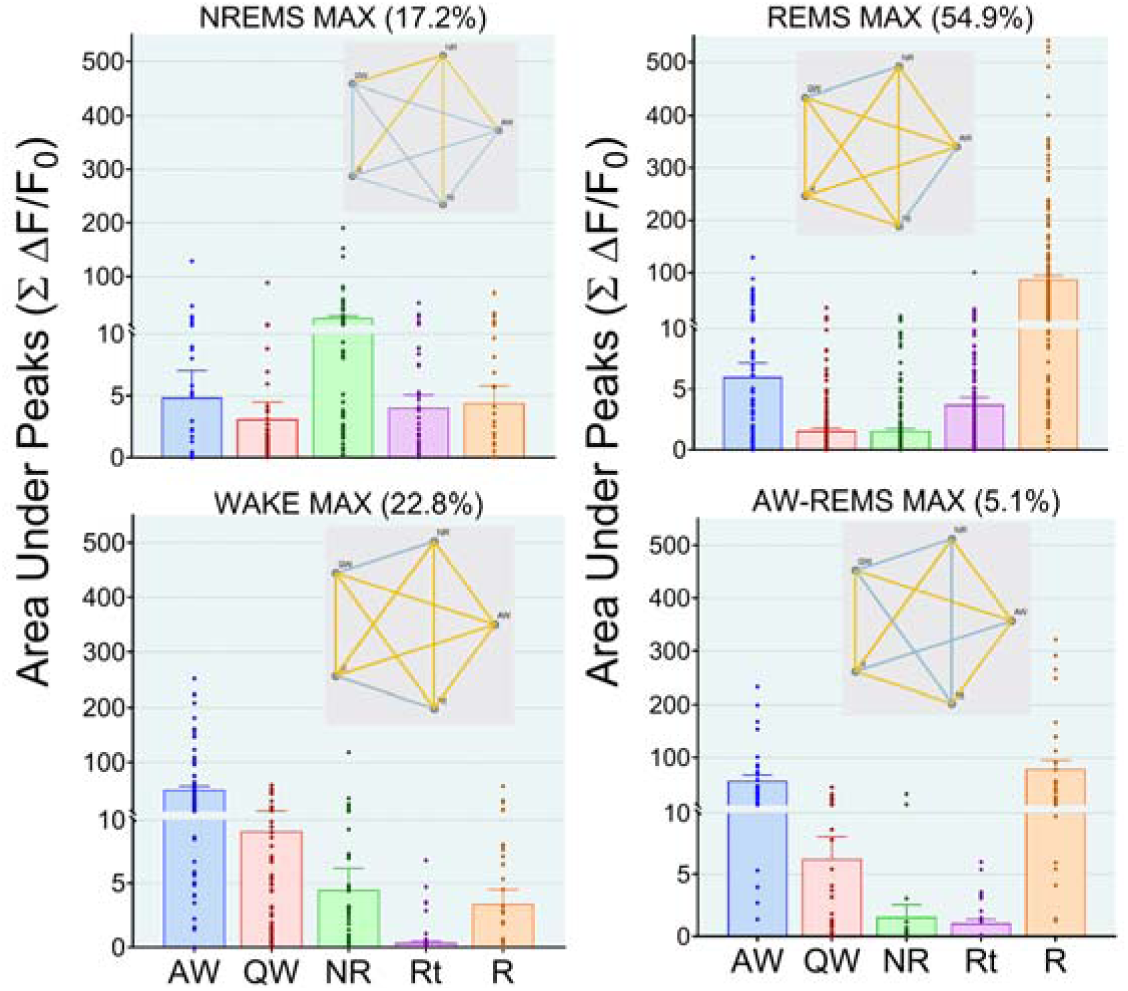
Fluorescence activity profiles of 372 vGAT neurons recorded in the ZI-dLH area in relation to vigilance states. 72% of the vGAT neurons were most active during sleep (NREM and REM). Pentagrams indicate statistical difference according to a general linear mixed model. All values were plotted as individual stacked dots. Bar height indicate the mean while error bars are +SEM. AW=active wake; QW=quiet wake: NR=non-REM sleep; Rt=transition to REM sleep; R=REM sleep.

The wake-max group (22.8%; n=85) were significantly more fluorescent when the mice were behaviorally active (grooming, walking, rearing) compared to periods of quiet wake, NREM and REM sleep. (F[4,305]=14.2, P=0.0001). During active behaviors the wake-max neurons had five-fold more activity than that during quiet wake.

Fluorescence in the NREM-max (17.2%; n=64) group was significantly higher compared to the other vigilance states (F[4,295]= 6.7; P=0.0001). On average, the fluorescence in the NREM-max neurons was twice compared to the other states. The fluorescence during the other vigilance states was uniformly low and not significantly different from each other.

The fourth group comprised of neurons that were fluorescent in both active waking and REM sleep (AW-REM-max; 5.1%; n=19). In this group the fluorescence during active waking and REM sleep was significantly higher compared to the other states.

### The time-course of activity during transition between states

Figure 3 summarizes the time course of the change in the fluorescence signal in the NREM-max (n=64), REM-max (n=204) and wake-max (n=85) groups. These three groups of GABA neurons represent 94.9% of all the neurons recorded in the study. To plot the time-course the activity of individual neurons was aligned with respect to the onset and offset of each vigilance state.

**Figure 3.**
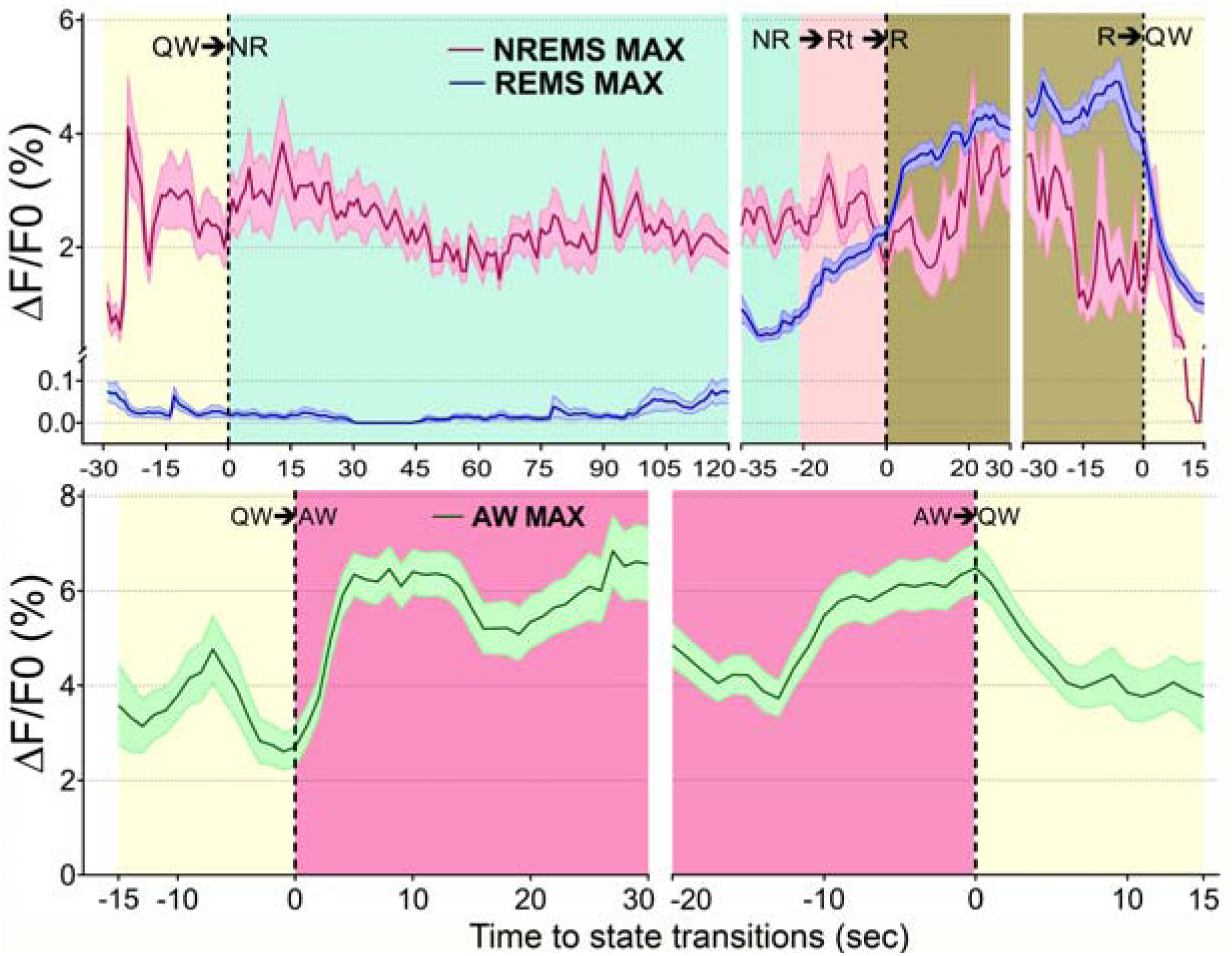
Time course of change in Ca2+fluorescence in vGAT neurons in the ZI-dLH during the transitions between sleep-wake states. The top panel depicts the time-course of activity in the NREM-max (64 neurons) and REM-max (204 neurons) groups while the lower panel depicts time-course of activity in the wake-max (85 neurons) group. The change in fluorescence was aligned to the transition point between states. Solid lines represent the mean fluorescence (ΔF/F0), while shaded bands indicate SEM. Notice that in the NREM-max neurons fluorescence increased 20s before the onset of NREM sleep and it remained elevated throughout NREM and REM sleep. In the REM-max group fluorescence was negligible at the start of NREM sleep, but significantly increased 35s before REM sleep onset and was elevated throughout REM sleep. In the wake-max neurons fluorescence occurred after the onset of behavioral activity and the fluorescence diminished only after the activity had ended.

The fluorescence in the neurons in the NREM-max group began to increase 20 sec prior to the onset of NREM sleep and the fluorescence remained high even during REM sleep. This suggests that these neurons may be instrumental in triggering both the behavioral and electrophysiological indices of sleep by actively suppressing the activity of wake-max arousal neurons.

The mean fluorescence in the REM-max group was low to negligible at the start of sleep but it increased sharply 35s before the onset of REM sleep. The fluorescence continued unabated during REM sleep and diminished only once the REM sleep episode had ended. It is important to note that the end of REM sleep is identified by the sudden re-emergence of muscle tone and behavioral activity. However, fluorescence did not increase indicating that the REM-max neurons are separate from the wake-max neurons.

In the wake-MAX group the transitions between AW↔QW were plotted (Figure 3). Activity in the wake-MAX neurons increased after the emergence of behavioral activity and remained high as long as the mice were behaviorally active (grooming, walking, rearing). Mice did not display feeding or drinking during our imaging sessions. Fluorescence subsided once the mice were quietly awake. Thus, the wake-max neurons did not anticipate changes in behavior.

## Discussion

The primary finding of our study was that 72% of the vGAT neurons in vZI/dLH were most active during sleep (NREM and REM sleep). These neurons increased their activity before the onset of sleep and continued to be active throughout the sleep episode. The heavy bias in activity of the vGAT neurons would initiate and maintain sleep serving as a counterweight to the activity of the arousal neurons. Indeed, GABA is the primary inhibitory neurotransmitter in the brain, and the most widely prescribed hypnotics induce sleep by enhancing the GABAergic inhibition across the entire CNS (Gottesmann, 2002).

### NREM-Max and REM-Max neurons

Another group has recorded activity of neurons in the same hypothalamic region as our study (Lee et al., 2005; Hassani et al., 2009; Hassani et al., 2010). They used the juxtacellular electrophysiological method to record activity of neurons in head-restrained rats (n=104 neurons in 77 rats) (Hassani et al., Eur J Neurosci, 2010). They found that 53% were sleep-max (n=55 of 104; NREM and REM-max), 18% wake-max (n=19 of 104), 17% wake-REM-max (n=18 of 104) and 11% of neurons that showed no change related to sleep-wake states (n=12 of 104). They then used post-mortem immunohistochemistry to confirm the phenotype identity of the neurons that were recorded based on intracellular-labeling of neurobiotin. They reported finding ten neurons that were positive for vGAT but negative for melanin concentrating hormone, and these were NREM-max (n=2) and REM-max (n=8). The two neurons that were NREM-max increased activity during the transition to NREM sleep. Our results are very similar to their results in that we both found NREM-max vGAT neurons that anticipated sleep onset (QW→NR), and were active during NREM sleep. The increase in action potentials during sleep in their study is similar to the increase in fluorescence that we measured using the genetically encoded Ca^2+^ sensor. The NREM-max group is the only group where the fluorescence increases with slow-wave EEG activity, a hallmark feature of the EEG that denotes NREM sleep.

### Wake MAX neurons

We found that a quarter of the recorded vGAT neurons in the ZI/dLH were wake-max while 5% were wake-REM max. These types of neurons were not recorded in the electrophysiology study but that is because in that study the rats were not freely-behaving. We suggest that allowing animals to move freely is critical to determine activity patterns of neurons. We observed that the wake MAX neurons became most active during grooming, walking, or rearing. We could not observe activity during feeding as the mice did not eat during the recording session.

### Activating vGAT-ZI neurons induces sleep

Clusters of vGAT neurons can colocalize other markers (calbindin, calretinin, galanin, LHx6, neuropeptide Y, parvalbumin, somatostatin) (Hernandez et al., 2015) (Qualls-Creekmore et al., 2017). Chemogenetic activation of the vGAT ZI cluster that coexpresses LHx6, the developmental gene transcriptional repressor, induces both NREM and REM sleep (Liu et al., 2017). The LHX6+neurons represent 45% of all vGAT+neurons within the vZI area, receive inputs from many arousal neurons and project to the orexin and GABA neurons residing in the vLH/PeF region. Indirect readout of their activity using c-FOS suggested the LHX6+vZI GABAergic neurons were most active during the lights-on phase which is the normal sleeping time in mice (Liu et al., 2017). These neurons also showed increased c-FOS in response to increased sleep pressure and sleep rebound. Most importantly, their study found that selective pharmacogenetic activation of the LHX6+vZI GABAergic neurons significantly enhanced both NREM and REM sleep. In contrast, selective inhibition of the LHx6 neurons significantly reduced both types of sleep. Thus, the selective manipulation of the spontaneous activity of the vZI GABAergic neurons mechanistically supports the role of the GABA neurons in the ZI/LH in inducing both NREM and REM sleep.

While the GABA neurons in the ZI-LH induce sleep there are other clusters of GABA neurons located ventrally and medially that induce waking and feeding behavior (Jennings et al., 2015). Pharmacogenetic activation of the ventral vGAT neurons increased waking, in particular feeding behavior (Venner et al., 2016). Interestingly, other active behaviors like climbing and grooming were strongly inhibited. Another group used optogenetics to stimulate the ventral vGAT neurons and found that it also increased waking (Herrera et al., 2016). Thus, although the ZI-LH area serves as an integrative node for regulation of wide variety of behaviors (Urbain and Deschenes, 2007; Wang et al., 2020), clusters of the GABA neurons can drive sleep.

## Acknowledgements

PJS was supported in part by the Department of Veterans Affairs, Veterans Health Administration, Office of Research Development (BLR&D; BX000798; BX004216), and NIH grants NS052287 and NS079940.

